# MUC5B and MUC5AC function in combination to regulate mucociliary transport on human airway epithelium

**DOI:** 10.64898/2026.09.08.750178

**Authors:** Sahana Kumar, Allison Boboltz, Gregg Duncan

## Abstract

Muco-obstructive lung diseases are characterized by impaired airway clearance and altered mucin composition. MUC5B and MUC5AC are the primary gel-forming mucins in airway mucus, yet how their relative abundance influences the physical and functional properties of mucus remains poorly understood. Here, we investigated how compositional variations in MUC5B and MUC5AC within mucus impact mucociliary transport. Mucus enriched in either MUC5B or MUC5AC was generated using human airway epithelial models depleted for each mucin via CRISPR/Cas9-targeted knockout. Defined mixtures of these mucins were generated at physiologically relevant ratios, for assessment of their microrheological properties and mucociliary transport behavior in differentiated primary human airway epithelial tissue cultures. We found that increasing MUC5AC content reduced network pore size and increased microviscosity, concomitant with reduced mucociliary transport, demonstrating that mucin composition alters the biophysical and functional properties of the mucus barrier. In normal airway tissue cultures with MUC5B-predominant mucus, overlay of MUC5AC on the apical surface impaired mucociliary transport, whereas MUC5B had minimal effect. Conversely, MUC5B supplementation uniquely improved mucociliary transport in IL-13 stimulated cultures exhibiting mucostasis, whereas additional MUC5AC did not alter transport. Together, these findings demonstrate that the ratio of MUC5B to MUC5AC can shape mucus organization at the microscale, which in turn governs mucociliary transport at the tissue scale.

## Introduction

The continuous production of mucus is crucial for protecting the airways and maintaining pulmonary health. Healthy airways are characterized by mucus secretions with low viscoelasticity that can be effectively cleared through mucociliary transport mechanisms.^1,2^ Chronic airway diseases can result from impaired mucociliary clearance (MCC) due to thick, hyper-concentrated mucus that is difficult to clear.^3,4^ In diseases such as asthma, chronic obstructive pulmonary disease (COPD), and cystic fibrosis (CF), pathological mucus results from changes in mucin composition and overall mucus amounts through the effects of mucin hypersecretion and/or airway dehydration.^5–7^ These changes hinder the clearance of mucus from the airways, resulting in mucus buildup, airway obstruction, and persistent inflammation creating an environment conducive to respiratory infections.^4,8,9^ Mucus is composed of extensively glycosylated gel-forming mucins, mucin 5B (MUC5B), and mucin 5AC (MUC5AC) that form a mucus gel network through disulfide bonds and electrostatic interactions.^10^ The resulting gel functions as a barrier to external substances and pathogens.^11–13^ Typically, in healthy airways, MUC5B is the predominant secreted mucin, and in comparison, MUC5AC is expressed at lower levels.^14,15^ However, in respiratory diseases, significant changes in mucin composition occur, resulting in mucus dysfunction and impaired airway clearance.

Based on biochemical analysis of sputum from asthmatics, there is a shift in the predominant mucin from MUC5B to MUC5AC.^16–19^ Studies conducted in murine and airway epithelial cell culture models of asthma have demonstrated that MUC5AC expression is upregulated in response to type 2 inflammation mediated by IL-13.^20–22^ Further work established IL-13 mediated overexpression of MUC5AC impairs mucus transport through tethering to the epithelium and enhancements in mucus viscoelasticity which contribute to airway obstruction in asthma.^23–25^ Mucin composition is also altered in COPD and CF, although reports differ as to whether MUC5B or MUC5AC predominates.^26–28^ Studies in sputum samples collected from human CF patients observed an overall increase in both MUC5B and MUC5AC leading to substantially higher total solids content.^29,30^ Increases in total mucin concentration or solids content of mucus can dehydrate the periciliary layer which compromises mucociliary transport in chronic lung diseases, particularly in CF.^31,32^ Together, these findings indicate alterations in mucin production and changes in mucus viscoelasticity and concentration drive airway clearance dysfunction in muco-obstructive lung diseases.

Investigations into the function of each mucin in the airway has revealed their unique roles in host defense. MUC5B is essential for functional MCC, with Muc5b (murine analog to MUC5B) deficiency in mice resulting in defective MCC leading to chronic bacterial infections and immune dysregulation.^33,34^ Similarly, recent studies in humans with congenital absence of MUC5B have shown impaired MCC.^35^ We have also shown in our prior work that the lack of MUC5B expression renders mucus immobile while the lack of MUC5AC expression impairs global coordination of mucus transport in human airway tissue cultures.^36^ However, there is still a limited understanding of how changes in relative abundance of MUC5B and MUC5AC, in combination, impacts the biological function of mucus as a protective barrier in the lung. At the biomolecular scale, MUC5B and MUC5AC exhibit distinct architectures and spatial organization within airway mucus. MUC5B predominantly forms long, bundled strands, whereas MUC5AC forms more highly branched, thread-like structures.^37,38^ Consistent with these differences, immunostaining of sputum and histological sections from individuals with asthma has revealed spatially distinct MUC5B-and MUC5AC-rich domains, suggesting that the two mucins can segregate within the mucus network.^23,39^ However, it has also been established MUC5B and MUC5AC can be co-packaged within the same secretory granules, indicating that these mucins are released together into the airway lumen.^40^ The blending of structurally distinct biopolymers can produce rheological properties that are not simply the sum of their parts, as inter- and intra-molecular interactions can drive complex network organization that generates emergent material behavior.^41–43^ For example, recent work has shown the blending of MUC5B and MUC5AC derived from porcine tissue substantially alters mucin chain reptation dynamics, rheological properties, and barrier function towards small molecule diffusion.^44^ Thus, the coexistence of structurally distinct mucins such as MUC5B and MUC5AC may impart physical properties to airway mucus that cannot readily be predicted from the properties of either mucin alone.

Herein, we examine the collective influence of MUC5B and MUC5AC on the physical nature and transport properties of airway mucus. For these studies, we cultured human airway epithelial (HAE) cells cultured at the air-liquid interface (ALI) which secrete their own mucus, with properties comparable to native airway secretions.^45–47^ However, we are unable to directly control and systematically vary MUC5B and MUC5AC expression using the standard HAE culture system. To address this, we used a previously established a BCi-NS1.1 HAE cell line with targeted knockout (KO) of either MUC5B or MUC5AC as a source of each individual mucin^333436^.^36^ Using this model, we collected mucus rich in either MUC5B or MUC5AC, respectively, and systematically varied the ratio of MUC5B to MUC5AC to investigate how changes in mucin composition affect mucus biophysical properties and mucociliary transport. Further, we evaluated mucociliary transport after exogenously manipulating MUC5B and MUC5AC in primary HAE cultures under basal conditions as well as asthma-like conditions with IL-13 stimulated MUC5AC hypersecretion. Collectively, our studies show MUC5B and MUC5AC contribute to mucociliary function in a synergistic manner that may either promote or impair airway clearance.

## Results

### Characterization of secreted mucin glycoproteins in MUC5B- and MUC5AC-deficient human airway epithelial cell culture models

As noted, we used BCi-NS1.1 cells, an immortalized airway epithelial basal cell line genetically engineered to achieve knockout (KO) of either MUC5AC or MUC5B.^36^ When cultured at an air-liquid interface (ALI), these cultures develop into functional muco-ciliated airway epithelium. We have previously established via Western blot and more recently via proteomic analysis that MUC5B and MUC5AC KO cultures showed selective depletion of the corresponding secreted mucin.^36,48^ To further characterize the knockout (KO) cultures, we performed lectin staining using wheat germ agglutinin (WGA) and Ulex europaeus agglutinin 1 (UEA1), which preferentially stain MUC5B and MUC5AC, respectively, due to their unique glycosylation profiles.^37,49^ WGA fluorescence intensity was increased in MUC5AC KO cultures (p=0.62), while UEA1 significantly increased in MUC5B KO cultures (p<0.0001), as shown in the representative fluorescence images (**Figure 1A**) and the quantified fluorescence integrated densities (**Figure S1**). Mucin expression was comparable across mucus collected from the KO cultures based upon fluorometric assessment of O-linked glycoprotein concentration (**Figure 1B**). To test whether exogenously added mucin could integrate within endogenously secreted mucus in HAE cultures, we supplemented MUC5AC KO and MUC5B KO cultures with 20 µl of exogenous MUC5AC-rich and MUC5B-rich mucus, respectively, and allowed the cultures to equilibrate for 1 hour. Treated cultures were then stained with WGA and UEA1 to determine whether mucociliary transport enabled spreading of exogenous mucin on the airway surface. After supplementation, changes in staining patterns with WGA and UEA1 were indicative of increased MUC5B and MUC5AC on the surface with spatial variation in their spread and qualitatively, appear to mix upon exogenous addition (**Figure 1C**).

**Figure 1.**
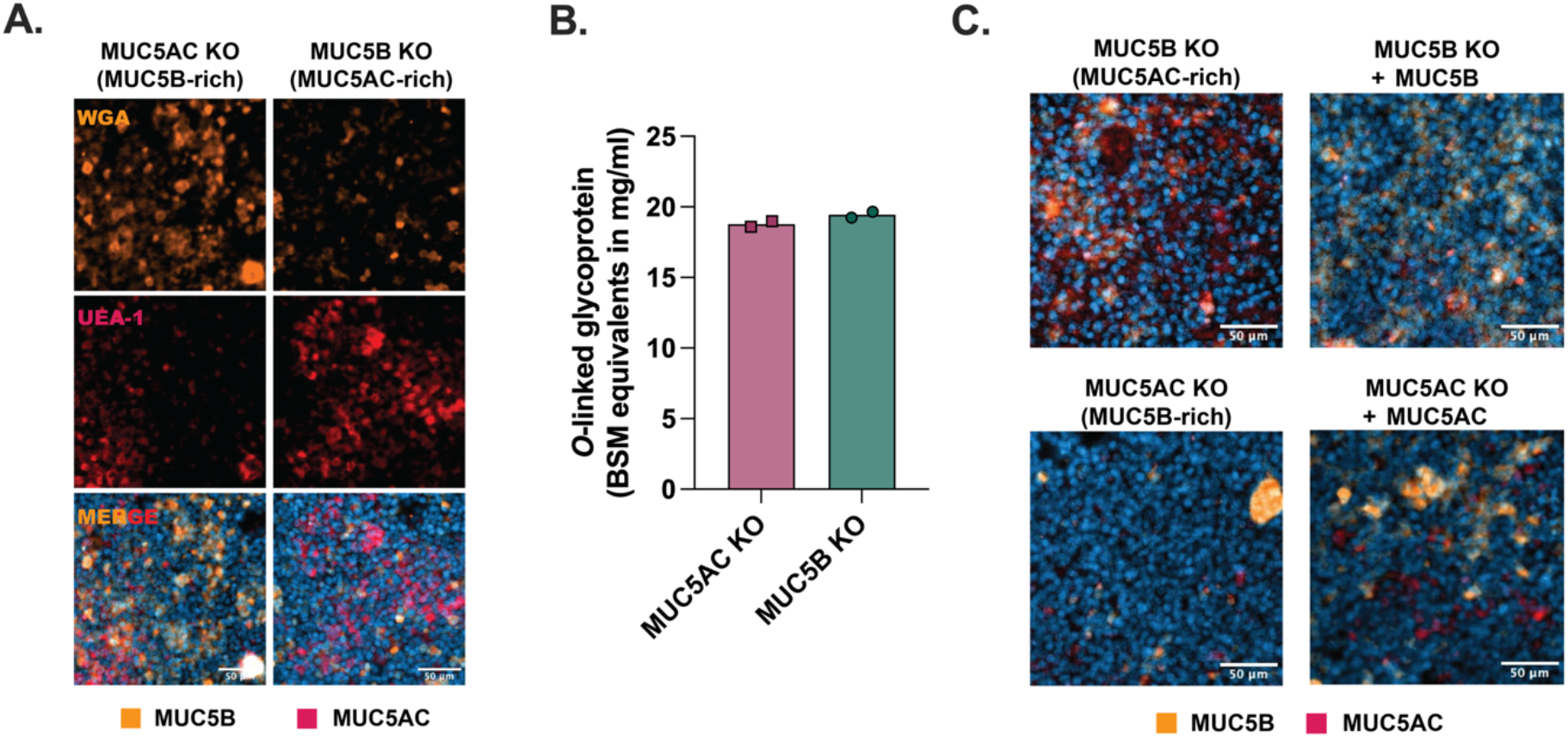
Characterization of secreted mucin glycoproteins in MUC5B- and MUC5AC-deficient HAE cultures. (**A**) Lectin staining of MUC5B and MUC5AC KO HAE cultures with WGA (orange) and UEA-1 (red). Scale bar = 50 µm (**B**) Fluorometric assessment of *O*-linked glycoprotein content as compared to bovine submaxillary mucin (BSM) as a standard. (**C**) Fluorescence images of lectin staining of MUC5B and MUC5AC KO cultures before and after supplementation with MUC5B and MUC5AC, respectively. Scale Bar = 50 µm.

### Microrheology of mucus with varying MUC5B:MUC5AC ratios

We generated stocks of MUC5B-rich and MUC5AC-rich mucus secretions from differentiated KO HAE cultures, measured their protein concentrations using the BCA assay, mixed the mucins at different mass ratios for subsequent analysis (**Figure 2A**). Based upon the production of secreted mucins in health and disease, we systematically varied the MUC5B:MUC5AC ratio in mucus gels as follows: (i) 75:25, (ii) 50:50 and (iii) 25:75. Gels prepared at these ratios were stained with WGA and Jacalin to visualize MUC5B and MUC5AC, respectively (**Figure S2**). To assess the effects of altered mucin composition on mucus gel properties, we used a tabletop microviscometer to measure apparent viscosity, and multiple particle tracking (MPT) to probe microrheology in gels with varying MUC5B:MUC5AC ratios. Representative particle trajectories illustrate the markedly reduced diffusion in gels containing higher MUC5AC content (**Figure 2B**). MPT analysis revealed that increasing MUC5AC content restricted the diffusion of fluorescently labeled 100-nm PEGylated nanoparticles as compared to mucus harvested from NHBE cultures (**Figure 2C**). Consistent with this observation, mean squared displacement over 1 s (MSD_1s_) was highest in 75% MUC5B/25% MUC5AC gels and significantly reduced in both the 50% MUC5B/MUC5AC and 25% MUC5B/75% MUC5AC gels (**Figure 2D, Figure S3A**). The reduced MSD indicates a more restrictive gel network, consistent with median pore size measurements derived from the MSD data (**Figure S3B**). Similarly, MPT-derived microviscosity increased with MUC5AC content, with comparable median values in the 50% and 25% MUC5B gels (∼3.2–3.9 mPa·s) that were significantly higher than that of the 75% MUC5B/25% MUC5AC gels (2.6 mPa·s) (**Figure S3C**). Using the microviscometer, apparent viscosity increased with MUC5AC content, with similar mean viscosities for the 50% MUC5B/MUC5AC and 25% MUC5B/75% MUC5AC gels (12.91 and 12.57 mPa·s, respectively), both of which were higher than that of the 75% MUC5B/25% MUC5AC gels (10.03 mPa·s) (**Figure 2C**).

**Figure 2.**
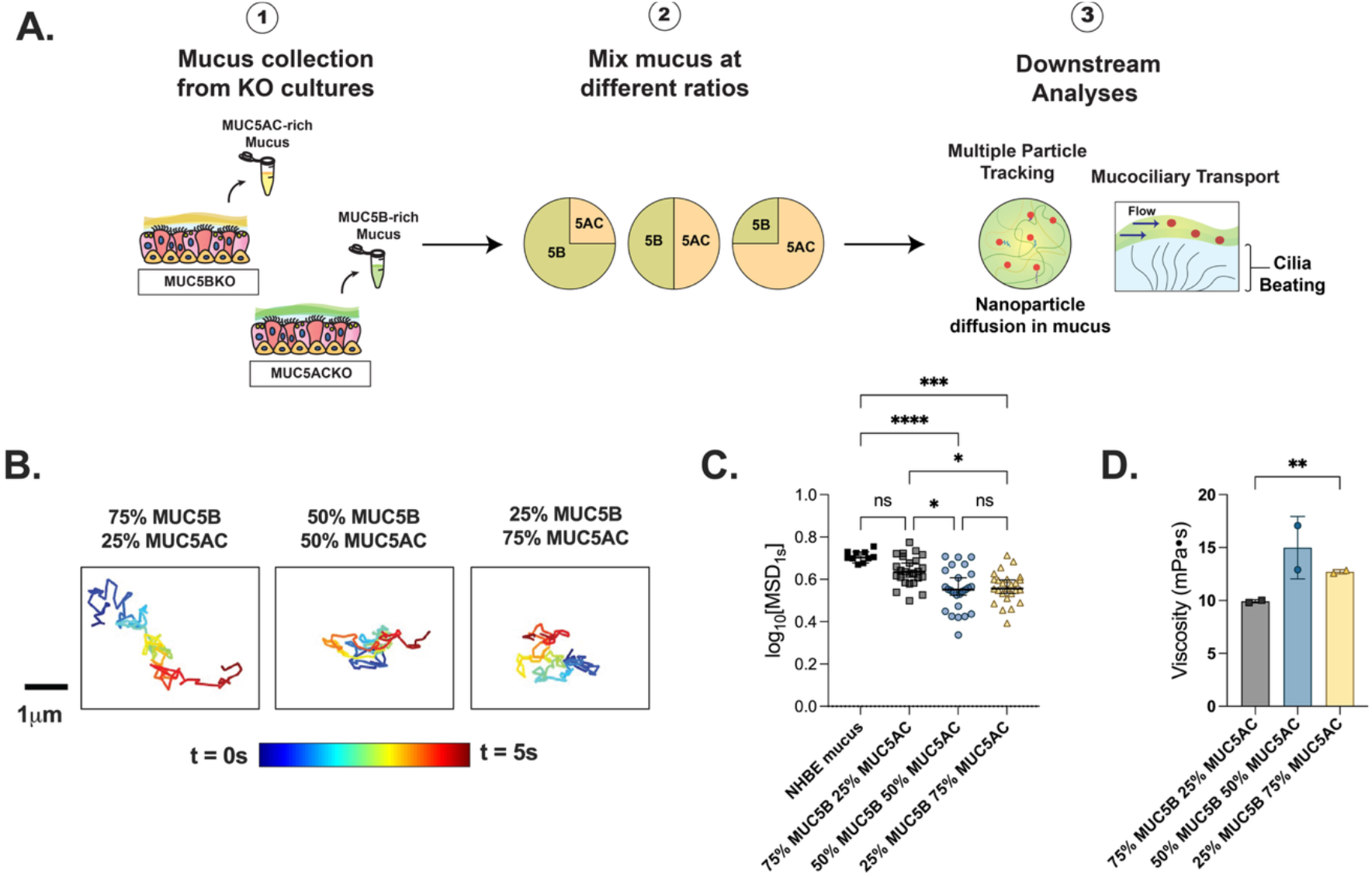
Microrheological properties of mucus gels with varying MUC5B and MUC5AC ratios. (**A**) Schematic illustration of preparing mucus gels with varying mucin composition. Mucus was exogenously collected from MUC5B & MUC5AC KO cultures. MUC5B and MUC5AC were mixed at varying ratios, and the resulting gels were used for downstream analyses. (**B**) Representative trajectories of 100nm muco-inert nanoparticle diffusion in mucus gels with MUC5B:MUC5AC ratios of 75:25, 50:50, 25: 75 (**C**) Scatter plot of measured median log10 [MSD_1s_] of 100nm NPs in mucus gels with varying mucin compositions. Each dot represents median measured MSD in individual videos taken from each biological replicate (n=3). Data sets statistically analyzed using Kruskal-Wallis test with Dunn’s test for multiple comparisons: *\*p*<0.05, \*\*\*\**p* < 0.0001. (**D**) Bar graph of measured apparent viscosities from a tabletop viscometer of mucus gels with varying mucus compositions. *\*\*p*<0.01 by Brown-Forsythe and Welch ANOVA.

### Transplantation of mucus gels with varying MUC5B:MUC5AC ratios in NHBE cultures and its impact on transport behavior

To evaluate the impact of varying mucin composition on MCT, we transplanted MUC5B/MUC5AC mucus gels onto freshly washed primary normal human bronchiolar epithelial cells (NHBEs) from healthy donors grown at an air-liquid interface (ALI) and allowed the cultures to equilibrate for 1 hour. We measured MCT rates by tracking 2 µm fluorescent beads added to the apical surface using video microscopy (**Figure 3A)**. We evaluated how transplanting MUC5B, MUC5AC, and mixtures of these gels affected MCT rates in NHBEs. For NHBE cultures treated with monomucin (100% MUC5B or MUC5AC) mucus, representative bead trajectories and median velocities showed no significant difference in transport rates between NHBE coated with MUC5B-rich and MUC5AC-rich gels alone (**Figure 3B, 3C**), and ciliary beat frequency was also comparable between groups (**Figure 3D**).

**Figure 3.**
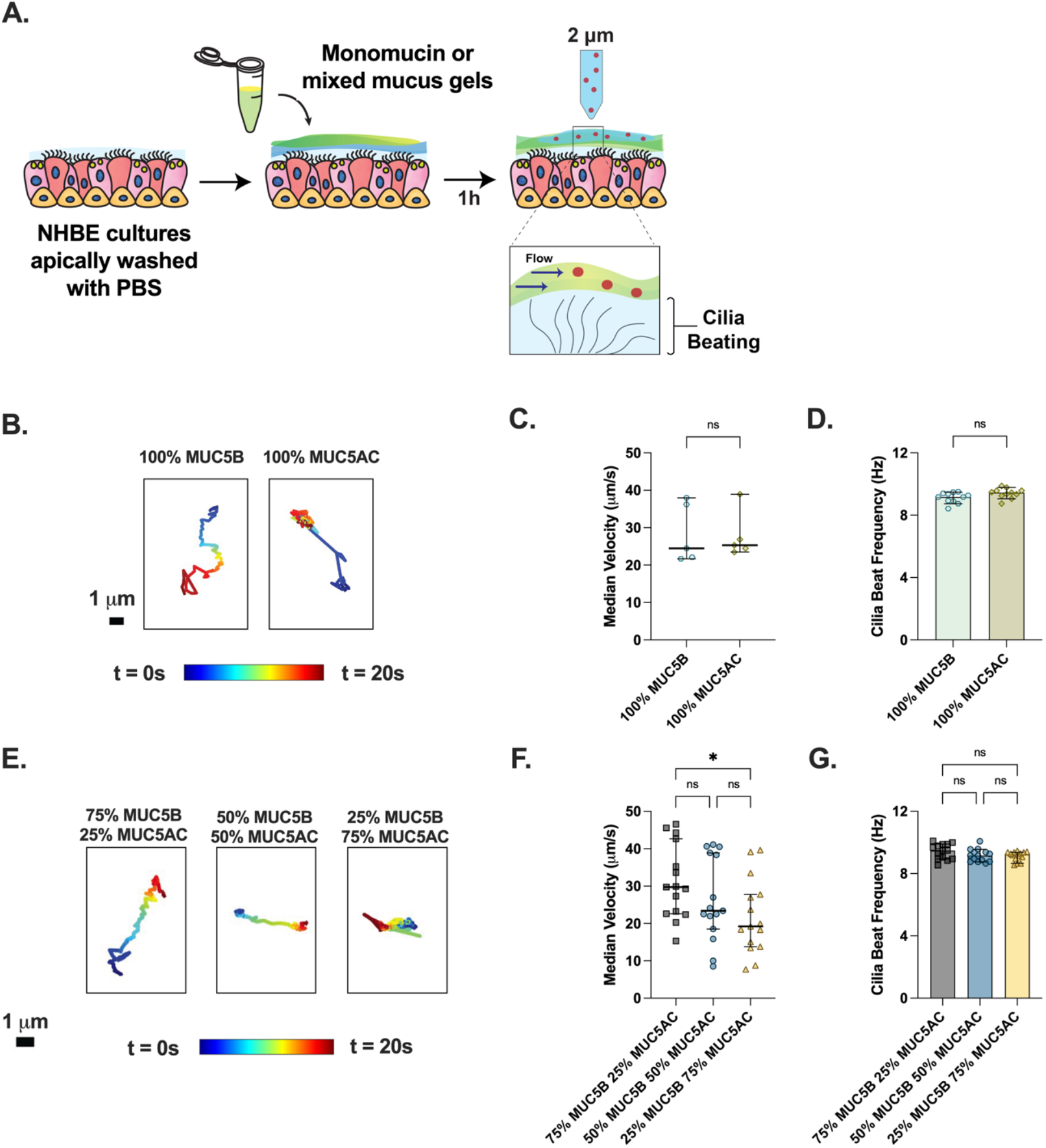
Mucociliary transport in differentiated primary NHBEs transplanted with mucus gels with varying mucin compositions. (**A**) Schematic illustration of experiment design. HAE cultures are washed apically with PBS and treated apically with monomucin (MUC5B, MUC5AC) or mixed MUC5B/MUC5AC gels with varied composition. After 1 h of equilibration, 2 μm beads are applied to the apical surface, and the cultures are imaged to track bead movement. **(B)** Representative trajectories of 2 µm beads in washed NHBEs supplemented with 100% MUC5B or 100% MUC5AC, respectively. (**C**) Scatter plot representing median velocities (µm/s) of beads in each video in NHBEs supplemented with MUC5B & MUC5AC. (**D**) Bar graph showing ciliary beat frequencies in NHBEs supplemented with MUC5B & MUC5AC. Each dot represents data from 1 video (n=3). Data sets in C,D statistically analyzed by two-tailed t-test. (**E**) Representative trajectories of 2 µm beads in washed NHBEs supplemented with mucus gels with 75% MUC5B/25% MUC5AC, 50% MUC5B/50% MUC5AC, and 25% MUC5B/75% MUC5AC, respectively. (**F**) Scatter plot representing median velocities (µm/s) of beads in each video in NHBEs supplemented with mixed mucus gels. (**G**) Bar graph showing ciliary beat frequencies in NHBEs supplemented with mixed mucus gels. Each dot represents data from 1 video (n≥3). Data sets in F,G statistically analyzed by one-way ANOVA (*\*p*<0.05).

In contrast, representative bead trajectories from the mixed mucus gels showed that beads traversed shorter distances in gels with higher MUC5AC content (50% and 75% MUC5AC) than in gels with higher MUC5B content (75% MUC5B) (**Figure 3E**). Quantification of MCT rates in NHBEs transplanted with these mixed mucus gels showed that velocities were highest in 75% MUC5B/25% MUC5AC gels (29.71 µm/s) (**Figure 3F)**, and decreased with increasing MUC5AC content, as seen in the 50% MUC5B/50% MUC5AC (23.38 µm/s and 25% MUC5B/75% MUC5AC (19.23 µm/s). MCT velocities in 75% MUC5B gels were approximately ∼1.5-fold higher than observed in 75% MUC5AC gels. Ciliary beat frequencies were comparable across all groups, indicating that the observed changes in MCT rates were not the result of changes in ciliary activity (**Figure 3G)**. We repeated the above studies in the immortalized cell line BCi-NS1.1 grown at ALI, following the same protocol outlined in **Figure 3A** yielding similar outcomes (**Figure S4**).

### Supplementation of normal and IL-13 stimulated airway cultures with exogenous MUC5B and MUC5AC and its differential impact on transport behavior

To determine the impact of exogenous mucin treatment on MCT, we supplemented NHBE with either MUC5B or MUC5AC at baseline or after IL-13 stimulated cultures to determine their impact on MCT (**Fig 4A**). Primary NHBEs can be stimulated to present an asthma-like phenotype *in vitro* by treating them with IL-13 at 10 ng/ml for 7 days, increasing MUC5AC secretion.^23–25^ Prior to supplementation with exogenous MUC5B or MUC5AC, IL-13 stimulated asthma-like cultures have reduced transport rates compared to healthy unstimulated (basal) NHBE cultures (**Figure 4B**). We observed that unstimulated NHBE cultures supplemented with MUC5AC exhibited significantly reduced mucociliary transport compared with cultures supplemented with MUC5B, which maintained transport comparable to untreated control cultures (**Figure 4C**). Supplementation with MUC5B significantly improved mucociliary transport in IL-13-stimulated NHBE cultures, whereas supplementation with MUC5AC did not significantly alter MCT as compared to untreated controls (**Figure 4D**). However, it should be noted differences in MCT were statistically not significant between MUC5B and MUC5AC-treated IL-13 NHBE cultures given the variation observed in these data sets. Similarly, we quantified changes in MCT following MUC5B and MUC5AC supplementation in KO cultures with MUC5B and MUC5AC-rich mucus production (**Figure S5A**). Consistent with our prior work,^36^ supplementation with MUC5B significantly improved MCT rates in MUC5B KO cultures, as shown by representative bead trajectories and median velocities (**Figure S5B**,**C**); velocities after supplementation were approximately 2.34-fold higher than before supplementation. MCT rates in MUC5AC KO (MUC5B-rich) cultures and in MUC5AC KO cultures supplemented with MUC5AC were comparable, indicating that supplementing the absent mucin did not significantly alter velocity in this case (**Figure S5D**,**E**) which is also consistent with our previous findings.^36^

**Figure 4.**
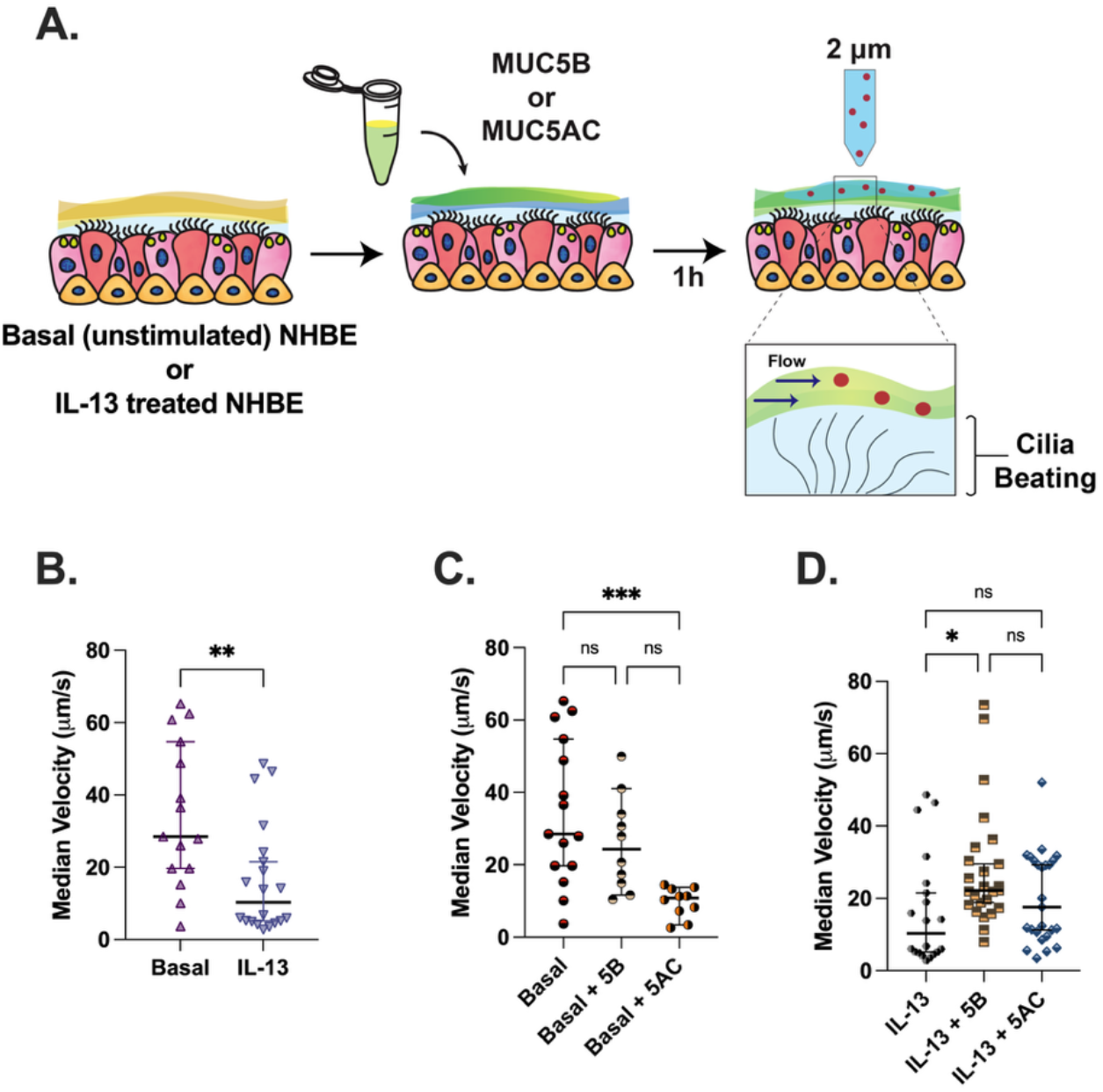
Mucociliary transport in untreated and IL-13 stimulated NHBEs supplemented with MUC5B or MUC5AC. (**A**) Schematic illustration of experiment design. Basal (unstimulated) or IL-13 stimulated cultures (10 ng/mL IL-13, 7 days) are supplemented with MUC5B or MUC5AC on the apical surface. After 1 h of equilibration, 2 µm beads are applied to the apical surface, and the cultures are imaged to track the bead movement. (**B**) Scatter plot representing median velocities (µm/s) of beads in healthy NHBE and NHBE treated with IL-13. Each dot represents data from 1 video (n≥3). (**C**) Scatter plot representing median velocities (µm/s) of beads in healthy NHBEs supplemented with MUC5B and MUC5AC. Each dot represents data from 1 video (n≥3). Data in B statistically analyzed by two-tailed t-test: \*\**p*<0.01. (**D**) Scatter plot representing median velocities (µm/s) of beads in NHBEs treated with IL-13. IL-13 supplemented with MUC5B and MUC5AC. Each dot represents data from 1 video (n≥3). Data sets in C,D statistically analyzed by ordinary one-way ANOVA: *\*p*<0.05, *\*\*\*p<*0.001.

## Discussion

Using our previously generated *in vitro* HAE models that are deficient in MUC5B or MUC5AC,^36^ we evaluated how the 2 major gel-forming airway mucins influence mucociliary clearance function when combined in physiologically relevant ratios. MUC5B-rich and MUC5AC-rich mucus was collected exogenously and mixed at defined ratios by mass to control their relative proportions across experiments. Lectin staining confirmed qualitatively that exogenously added mucins integrated with endogenously secreted mucus within HAE cultures. Although the resulting mucus appeared heterogeneous, we did not observe clear evidence of discrete phases, suggesting that MUC5B and MUC5AC coexist within a common mucus network. We recognize our studies may not reflect the organization of secreted mucins observed in disease, such as in asthma characterized with MUC5AC overproduction, where regions of the airway could produce mucus enriched in a disease-associated mucin subtype. We also evaluated the effect of these mucins when added exogenously in the context of normal and dysfunctional MCT. Collectively, these studies suggest that airway mucus function is governed not only by mucin abundance, but also by the relative composition of each constituent mucin.

As noted, previous studies have visualized mucin structures using scanning electron microscopy and observed that MUC5AC forms a branched network, while MUC5B forms linear, less branched structures that bundle.^37,38^ This motivated our studies to evaluate how mixing of these mucins with distinct structures may augment mucus gel properties. Previous work by our group and others has shown that MUC5AC forms a tighter mesh microstructure thought to significantly enhance mucus elasticity and adhesivity.^23,36,38^ Consistent with this, we find MUC5AC-predominant (≥50% MUC5AC) gels possess significant reductions in network pore size and increases in microviscosity as compared to MUC5B-predominant gels. This suggests the increasing prevalence of MUC5AC in airway mucus increases the viscoelastic properties which in turn influence mucus transport function. We note that viscoelastic properties of mucus contribute to MCT in parallel with other factors including concentrationdependent surface hydration and mucus-epithelial cell surface adhesion which may depend on both mucin concentration and composition.^31,50,51^

In our prior work, we found that each mucin had unique roles in supporting mucus transport where MUC5B enabled mobilization of mucus under ciliary action whereas MUC5AC promoted directional flow alignment.^36^ In addition, we showed addition of MUC5B to MUC5AC–secreting cultures normalized MCT which was further confirmed in this current work. However, it is unclear from this data alone if this improvement arose from reductions in mucus viscoelasticity and/or reversal of MUC5AC-mediated mucus gel tethering to the epithelium. Building on this work, we find here the transport of MUC5B and MUC5AC gels on the airway surface are similar in nature when added exogenously to primary HAE cultures. This outcome suggests MUC5AC-enriched gels are not innately less transportable than MUC5B. While we did not expect such similarity in transport in MUC5B and MUC5AC-coated NHBE cultures, we note our findings are similar to what was observed in a MUC5AC-overexpressing transgenic mouse model where no defects in MCT were observed despite the enrichment of MUC5AC in the airways.^52^ Moreover, it is possible these similarities in transport are due to both of the MUC5B-rich and MUC5AC-rich gel preparations being in a normal, well-hydrated state. It is very possible MUC5B and MUC5AC gels would possess distinct behavior in a more dehydrated and concentrated state. However based on the studies herein, MUC5AC-mediated impairment of MCT is presumably due to incomplete release and direct surface tethering of MUC5AC to the underlying epithelium. However, deficits in MCT were observed in a MUC5AC concentration-dependent manner for gels containing a mixture of MUC5B and MUC5AC. These data suggest the viscoelastic nature as well as interfacial interactions of mucus with the epithelium are distinct in mixtures of MUC5B and MUC5AC as compared to gels composed of each individually.

Our previous work demonstrated that exogenous supplementation of airway surface liquid with MUC5B or MUC5AC can alter mucus transport behavior.^36^ However, the extent and robustness of MCT in the BCi-NS1.1 HAE model does not match that observed in primary NHBE cultures. Further, it has yet to be determined if the beneficial effects of MUC5B under MUC5AC-dominant conditions extend to an IL-13-induced state of MUC5AC hypersecretion. We therefore examined the effects of exogenous MUC5B and MUC5AC on MCT in primary NHBE cultures under basal conditions and following IL-13 stimulation. MUC5AC supplementation impaired transport in healthy NHBE cultures with a baseline MUC5B-predominant secretion profile, whereas MUC5B had minimal effect. Conversely, MUC5B supplementation improved transport in IL-13-stimulated cultures, whereas additional MUC5AC did not influence transport. Notably, the observed effects occurred despite the substantial increases in total mucus load imposed by exogenous mucin supplementation, which would be expected to impede rather than enhance transport independent of the type of mucin added. Together, these findings demonstrate that mucus transport depends strongly on mucin composition. Even in the IL-13-induced pathological state, MUC5AC-associated MCT dysfunction was found to be, at least partially, reversible upon addition of MUC5B. Based on our studies with monomucin gel transport on NHBE cultures, it is likely the observed mucin-specific transport behavior is strongly influenced by interfacial interactions between mucus and the airway epithelium. Considering this, it would be important in future studies to evaluate alterations to the glycocalyx composition and tethered mucin expression profile under normal and IL-13 stimulated conditions where MUC5B and MUC5AC may possess differential adhesion to a healthy versus diseased airway epithelium.

We recognize several limitations of this work. First, our KO culture models while heavily enriched with MUC5B and MUC5AC are not entirely pure species based upon Western blot and proteomic analysis in our previous work.^36,48^ However, we did not purify the mucus from these cultures further as additional processing would disrupt their native biophysical characteristics. More in-depth characterization using bulk rheological methods may provide additional information directly applicable to the mechanical response of mucus gels under ciliary action. The primary mechanism(s) driving the emergent properties of MUC5B and MUC5AC are also not fully defined through this work. We propose several means by which MUC5B and MUC5AC may generate gels with unique transport properties through mucin-mucin intermolecular interactions and network assemblies. For example, the reductions in network pore size in MUC5AC-dominant gels could both lead to altered viscoelasticity as well as mucus-surface adhesion and anchoring. Moreover, differential glycosylation of MUC5B and MUC5AC could also be a contributing factor in the assembly and interfacial adhesion of mucus gels which is unfortunately beyond the scope of our current work. Future studies on the adhesion at airway mucosal interfaces for monomucin and mixed gels composed of MUC5B and MUC5AC may help elucidate its contributions to our findings here. Overall, these findings establish mucin composition as a determinant of airway mucus function and demonstrate that the relative abundance of MUC5B and MUC5AC can shape the physical and transport properties of mucus. More broadly, our findings provide a framework for understanding how changes in MUC5B and MUC5AC influence mucus function in airway disease.

## Materials & Methods

### Cell Culture

MUC5B and MUC5AC knockout (KO) cultures were generated as previously described^36^ in BCI-NS1.1 cells, a human immortalized basal cell line provided and characterized by Ronald Crystal’s group (Weill Cornell Medical College).^53^ Primary Normal Human Bronchiolar Epithelial (NHBE) cells were obtained from Lonza. The cultures were maintained at 37°C, 5% CO_2_ in a flask with PneumaCult-Ex Plus expansion media (STEMCELL Technologies). The cells were detached at 80% confluency using 0.05% Trypsin-EDTA for 5 minutes and seeded onto 12-mm transwell inserts (STEMCELL Technologies) coated with rat-tail collagen type I at 10,000 cells/cm^2^ in ExPlus medium supplemented to both the apical and basolateral compartments. After the cells reached 100% confluency, the media on the apical surface was removed, and the basolateral media was replaced with PneumaCult-ALI media (STEMCELL Technologies). The cells were induced to differentiate for 28 days, with media changed every other day.

### Mucus Collection and Mucus Gel Preparation

After the cells reached differentiation, mucus was collected from the apical surface of MUC5B and MUC5AC KO cultures twice a week. Sterile PBS was added to the apical surface for 30 min at 37°C and collected. The washings were passed through 100 kDa Amicon filters at 14,000 x g for 20 min and stored at -80°C for long-term storage prior to use. The protein in mucus samples was measured using the Pierce BCA Protein Assay kit (ThermoFisher Scientific) prior to mixing MUC5B and MUC5AC KO at 25:75, 50:50, and 75:25 ratios based on protein mass. Based upon comparison to bovine serum albumin (BSA) as a standard, total protein content ranged from ∼3–7 mg/mL in pooled mucus washings from our cultures. Mixtures of MUCB KO and MUC5AC KO culture-derived mucus gels were then prepared at a fixed total protein concentration (BSA-equivalent) of ∼3 mg/mL. Once prepared, the mucus gels were allowed to equilibrate for 24 hours at 4°C before biophysical characterization, transport, lectin, and immunostaining experiments.

### Particle Tracking Microrheology (PTM)

100nm carboxylate-modified polystyrene nanoparticles (ThermoFisher Scientific) were coated with 5 kDa methoxy-terminated polyethylene glycol (PEG) to ensure that these particles were muco-inert.^54^ The size and zeta potential of the nanoparticles were measured to ensure a near-neutral charge and approximately 100-150 nm size range, using NanoBrook Omni (Brookhaven Instruments). To image nanoparticle diffusion within mucus, 20 µl of mucus gels containing fixed total protein mass (∼60 µg, BSA-equivalents) were added to microscopy chambers made from vacuum-grease coated O-rings. To this, 1 µl of PEG-coated nanoparticles was added and allowed to equilibrate for 30 minutes at room temperature. Nanoparticle diffusion was measured by fluorescence video microscopy using a Zeiss 800 LSM microscope with a 63x water-immersion objective at a 33Hz frame rate for 10s. The data was analyzed based on a previously developed MATLAB algorithm.^55^ The mean-squared displacement (MSD) as a function of lag time was calculated for each particle. The viscoelastic properties of the mucus gels, such as pore size and microviscosity, were calculated from MSD based on the generalized Stokes-Einstein equation.

### Lectin Staining of HAE Cultures

The cultures were fixed with 100% methanol for 15 minutes at 4°C. The cultures were washed with PBS and blocked in carbo-free blocking buffer (Vector Laboratories) for 30 minutes at room temperature. The fluorescently labeled lectins Rhodamine-WGA, FITC-Jacalin, or Alexa Fluor 647-UEA1 were diluted to 1:1000 in PBS and incubated overnight at 4 °C with cultures. The cultures were washed twice with PBS and incubated with 1 µl/ml DAPI in PBS for 15 minutes at room temperature. These cultures were washed twice with PBS, mounted onto cover slides, and imaged using a Zeiss LSM 800 Microscope.

### Lectin Staining of Mucins

The mucus samples at different ratios were smeared onto glass slides and allowed to air-dry. The slides were fixed in 100% ice-cold methanol for 15 min at 4°C. The slides were washed with PBS for 5 mins at room temperature. The slides were blocked with a carbo-free blocking buffer (Vector Laboratories) for 30 mins at room temperature and washed with PBS for 5 mins. The slides were incubated with lectins, Rhodamine-WGA (Vector Laboratories) and Fluorescein-Jacalin (Vector Laboratories) at a 1:1000 dilution in PBS overnight at 4°C. The slides were washed twice with PBS for 5 min each and mounted with Vectashield mounting medium. The slides were imaged using the Zeiss LSM Microscope at a 10x objective.

### Mucociliary Transport and Ciliary Beat Frequency

To measure the transport of mucus gels containing defined ratios of MUC5B and MUC5AC, HAE cultures are washed, and immediately after, 20 µl pre-mixed mucus gels containing fixed total protein mass (∼60 µg, BSA-equivalents) are supplemented onto the apical surface of ALI cultures. The gels were allowed to equilibrate for 1 hour at 37°C, 5% CO_2_, after which 25 µl of 2 µm fluorescent microspheres (Invitrogen, diluted 1:2000 in PBS) was added apically and allowed to equilibrate for 15 mins at 37°C. Particle velocimetry measurements were then conducted to measure mucociliary transport (MCT). In each culture, videos from five different regions were recorded at 10x magnification at a frame rate of 10 Hz for 20 s and analyzed using a custom MATLAB algorithm.^56,57^ Ciliary beat frequency was analyzed based on 10 s videos at a frame rate of 50 Hz at 10x magnification from 3 random regions using brightfield. For assessment of MCT in KO cultures supplemented with the absent mucus, the cultures were washed 48 hours prior, and mucus was allowed to accumulate before supplementing with 20 µl of the knocked-out mucus. Cultures were then to equilibrated for 1 hour before adding 2 µm fluorescent microspheres to measure MCT.

## Acknowledgments

This project was funded by the Cystic Fibrosis Foundation (DUNCAN24G0), the National Institutes of Health (R01 HL160540, F31 HL176146), and University of Maryland Grand Challenges Research Grant (GC 18).

## Author contributions

S.K. and A.B. conceived, designed, and performed the research and led data analysis. G.A.D. conceived and designed experiments. S.K. and G.A.D. wrote the article. All authors reviewed and edited the article.

## References

1. Knowles, M. R. & Boucher, R. C. Mucus clearance as a primary innate defense mechanism for mammalian airways. J Clin Invest 109, 571–577 (2002).

2. Thornton, D. J. & Sheehan, J. K. From mucins to mucus: toward a more coherent understanding of this essential barrier. Proc Am Thorac Soc 1, 54–61 (2004).

3. Cone, R. A. Barrier properties of mucus. Advanced Drug Delivery Reviews 61, 75–85 (2009).

4. Fahy, J. V. & Dickey, B. F. Airway Mucus Function and Dysfunction. New England Journal of Medicine 363, 2233–2247 (2010).

5. Boucher, R. C. Muco-Obstructive Lung Diseases. New England Journal of Medicine 380, 1941–1953 (2019).

6. Evans, C. M., Kim, K., Tuvim, M. J. & Dickey, B. F. Mucus hypersecretion in asthma: causes and effects. Current Opinion in Pulmonary Medicine 15, 4–11 (2009).

7. Hill, D. B., Button, B., Rubinstein, M. & Boucher, R. C. Physiology and pathophysiology of human airway mucus. Physiological Reviews 102, 1757–1836 (2022).

8. Kuyper, L. M. et al. Characterization of airway plugging in fatal asthma. Am J Med 115, 6–11 (2003).

9. Boucher, R. C. On the Pathogenesis of Acute Exacerbations of Mucoobstructive Lung Diseases. Ann Am Thorac Soc 12 Suppl 2, S160–3 (2015).

10. Song, D., Cahn, D. & Duncan, G. A. Mucin Biopolymers and Their Barrier Function at Airway Surfaces. Langmuir 36, 12773–12783 (2020).

11. Wagner, C. E., Wheeler, K. M. & Ribbeck, K. Mucins and Their Role in Shaping the Functions of Mucus Barriers. Annu Rev Cell Dev Biol 34, 189–215 (2018).

12. Sheng, Y. H. & Hasnain, S. Z. Mucus and Mucins: The Underappreciated Host Defence System. Front. Cell. Infect. Microbiol. 12, 856962 (2022).

13. Witten, J., Samad, T. & Ribbeck, K. Selective permeability of mucus barriers. Current Opinion in Biotechnology 52, 124–133 (2018).

14. Okuda, K. et al. Localization of Secretory Mucins MUC5AC and MUC5B in Normal/Healthy Human Airways. Am J Respir Crit Care Med 199, 715–727 (2019).

15. Kesimer, M. et al. Airway Mucin Concentration as a Marker of Chronic Bronchitis. N Engl J Med 377, 911–922 (2017).

16. Nguyen, L. P. et al. Chronic exposure to beta-blockers attenuates inflammation and mucin content in a murine asthma model. Am J Respir Cell Mol Biol 38, 256–262 (2008).

17. Reader, J. R. et al. Pathogenesis of mucous cell metaplasia in a murine asthma model. Am J Pathol 162, 2069–2078 (2003).

18. Kirkham, S., Sheehan, J. K., Knight, D., Richardson, P. S. & Thornton, D. J. Heterogeneity of airways mucus: variations in the amounts and glycoforms of the major oligomeric mucins MUC5AC and MUC5B. Biochem J 361, 537–546 (2002).

19. Woodruff, P. G. et al. T-helper type 2-driven inflammation defines major subphenotypes of asthma. Am J Respir Crit Care Med 180, 388–395 (2009).

20. Kuperman, D. A. et al. Direct effects of interleukin-13 on epithelial cells cause airway hyperreactivity and mucus overproduction in asthma. Nat Med 8, 885–889 (2002).

21. Kuperman, D. A. et al. Dissecting asthma using focused transgenic modeling and functional genomics. J Allergy Clin Immunol 116, 305–311 (2005).

22. Zhen, G. et al. IL-13 and Epidermal Growth Factor Receptor Have Critical but Distinct Roles in Epithelial Cell Mucin Production. Am J Respir Cell Mol Biol 36, 244–253 (2007).

23. Bonser, L. R., Zlock, L., Finkbeiner, W. & Erle, D. J. Epithelial tethering of MUC5AC-rich mucus impairs mucociliary transport in asthma. Journal of Clinical Investigation 126, 2367–2371 (2016).

24. Kuyper, L. M. et al. Characterization of airway plugging in fatal asthma. Am J Med 115, 6–11 (2003).

25. Liegeois, M. A. et al. Cellular and molecular features of asthma mucus plugs provide clues about their formation and persistence. Journal of Clinical Investigation 135, e186889 (2025).

26. Radicioni, G. et al. Airway mucin MUC5AC and MUC5B concentrations and the initiation and progression of chronic obstructive pulmonary disease: an analysis of the SPIROMICS cohort. Lancet Respir Med 9, 1241–1254 (2021).

27. Caramori, G. et al. MUC5AC expression is increased in bronchial submucosal glands of stable COPD patients. Histopathology 55, 321–331 (2009).

28. Kirkham, S., Sheehan, J. K., Knight, D., Richardson, P. S. & Thornton, D. J. Heterogeneity of airways mucus: variations in the amounts and glycoforms of the major oligomeric mucins MUC5AC and MUC5B. Biochem J 361, 537–546 (2002).

29. Henke, M. O., John, G., Germann, M., Lindemann, H. & Rubin, B. K. MUC5AC and MUC5B mucins increase in cystic fibrosis airway secretions during pulmonary exacerbation. Am J Respir Crit Care Med 175, 816–821 (2007).

30. Niv, Y., Ho, S. B. & Rokkas, T. Mucin Secretion in Cystic Fibrosis: A Systematic Review. Dig Dis 39, 375–381 (2021).

31. Button, B. et al. A periciliary brush promotes the lung health by separating the mucus layer from airway epithelia. Science 337, 937–941 (2012).

32. Hill, D. B. et al. A Biophysical Basis for Mucus Solids Concentration as a Candidate Biomarker for Airways Disease. PLoS One 9, e87681 (2014).

33. Roy, M. G. et al. Muc5b is required for airway defence. Nature 505, 412–416 (2014).

34. Patil, N. et al. Dysregulated MUC5B and MUC5AC impair epithelial barrier function and alter granulocyte frequency and activation in the lung and distal sites. Mucosal Immunology 100385 (2026) doi:10.1016/j.mucimm.2026.100385.

35. Costain, G. et al. Hereditary Mucin Deficiency Caused by Biallelic Loss of Function of MUC5B. Am J Respir Crit Care Med 205, 761–768 (2022).

36. Song, D. et al. MUC5B mobilizes and MUC5AC spatially aligns mucociliary transport on human airway epithelium. Science Advances 8, eabq5049 (2022).

37. Ostedgaard, L. S. et al. Gel-forming mucins form distinct morphologic structures in airways. Proc. Natl. Acad. Sci. U.S.A. 114, 6842–6847 (2017).

38. Carpenter, J. et al. Assembly and organization of the N-terminal region of mucin MUC5AC: Indications for structural and functional distinction from MUC5B. Proceedings of the National Academy of Sciences 118, e2104490118 (2021).

39. Lachowicz-Scroggins, M. E. et al. Abnormalities in MUC5AC and MUC5B Protein in Airway Mucus in Asthma. Am J Respir Crit Care Med 194, 1296–1299 (2016).

40. Hoang, O. N. et al. Mucins MUC5AC and MUC5B Are Variably Packaged in the Same and in Separate Secretory Granules. American Journal of Respiratory and Critical Care Medicine 206, 1081–1095 (2022).

41. Frith, W. J. Mixed biopolymer aqueous solutions – phase behaviour and rheology. Advances in Colloid and Interface Science 161, 48–60 (2010).

42. Boyd, R. & Smith, G. Polymer Dynamics and Relaxation. (Cambridge University Press, 2007). doi:10.1017/CBO9780511600319.

43. Tucker Iii, C. L. & Moldenaers, P. M ICROSTRUCTURAL E VOLUTION IN P OLYMER B LENDS. Annu. Rev. Fluid Mech. 34, 177–210 (2002).

44. Henkel, F. & Lieleg, O. Foreign Mucins Alter the Properties of Reconstituted Gastric Mucus. Biomacromolecules 26, 2293–2303 (2025).

45. Kesimer, M. et al. Tracheobronchial air-liquid interface cell culture: a model for innate mucosal defense of the upper airways? Am J Physiol Lung Cell Mol Physiol 296, L92–L100 (2009).

46. Hill, D. B. & Button, B. Establishment of respiratory air-liquid interface cultures and their use in studying mucin production, secretion, and function. Methods Mol Biol 842, 245–258 (2012).

47. Kim, K. C. Biochemistry and pharmacology of mucin-like glycoproteins produced by cultured airway epithelial cells. Exp Lung Res 17, 533–545 (1991).

48. Corkran, M. et al. An Inverse Transwell Assay for Airway Mucus Barrier Function Reveals both Virus- and Mucin-Specific Impacts on Infection. Preprint at 10.64898/2026.07.19.739437 (2026).

49. García-Posadas, L. et al. Interaction of IFN-γ with cholinergic agonists to modulate rat and human goblet cell function. Mucosal Immunol 9, 206–217 (2016).

50. Button, B. et al. Roles of mucus adhesion and cohesion in cough clearance. Proceedings of the National Academy of Sciences 115, 12501–12506 (2018).

51. Abdullah, L. H. et al. Mucin Production and Hydration Responses to Mucopurulent Materials in Normal versus Cystic Fibrosis Airway Epithelia. Am J Respir Crit Care Med 197, 481–491 (2018).

52. Ehre, C. et al. Overexpressing mouse model demonstrates the protective role of Muc5ac in the lungs. Proceedings of the National Academy of Sciences 109, 16528–16533 (2012).

53. Walters, M. S. et al. Generation of a human airway epithelium derived basal cell line with multipotent differentiation capacity. Respir Res 14, 135 (2013).

54. Duncan, G. A. et al. Microstructural alterations of sputum in cystic fibrosis lung disease. JCI Insight 1, e88198 (2016).

55. Schuster, B. S., Ensign, L. M., Allan, D. B., Suk, J. S. & Hanes, J. Particle tracking in drug and gene delivery research: State-of-the-art applications and methods. Adv Drug Deliv Rev 91, 70–91 (2015).

56. Corkran, M., Boboltz, A., Duncan, G. A. & Scull, M. A. Methods for Discerning the Impact of Mucus on Host Defenses Against Viral Infection. Current Protocols 5, e70201 (2025).

57. Song, D. et al. Modeling Airway Dysfunction in Asthma Using Synthetic Mucus Biomaterials. ACS Biomater. Sci. Eng. 7, 2723–2733 (2021).

