## Supplementary Information for "MUC5B and MUC5AC function in combination to regulate mucociliary transport on human airway epithelium"

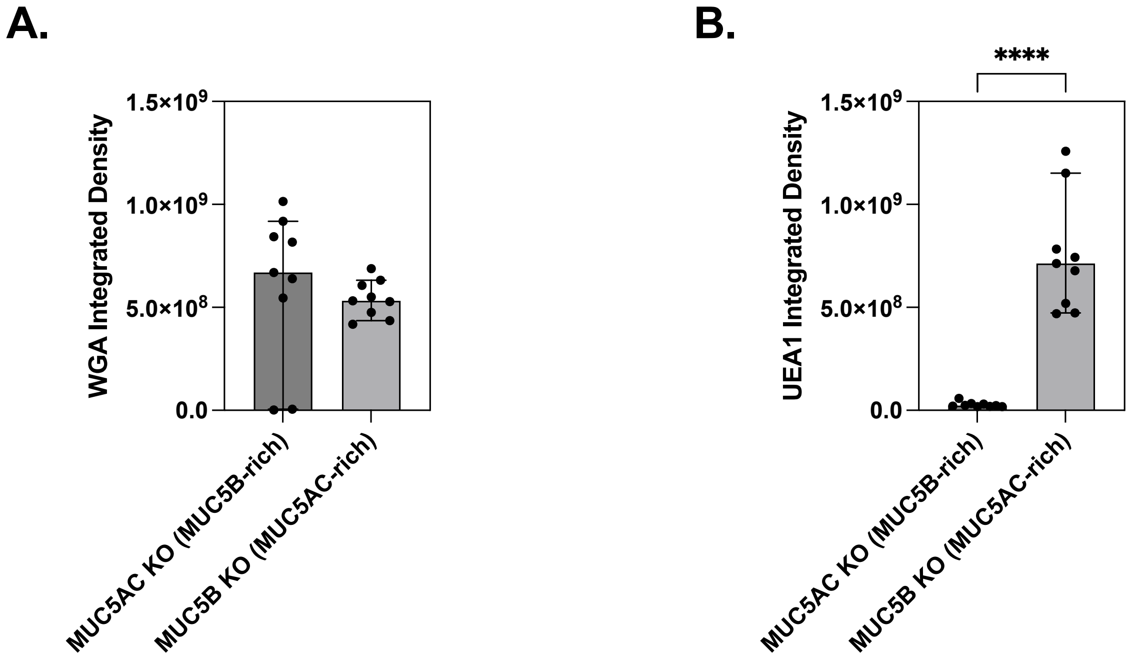


**Figure S1. Quantification of lectin staining** (**A**) WGA and (**B**) UEA1 of mucus from KO cultures.

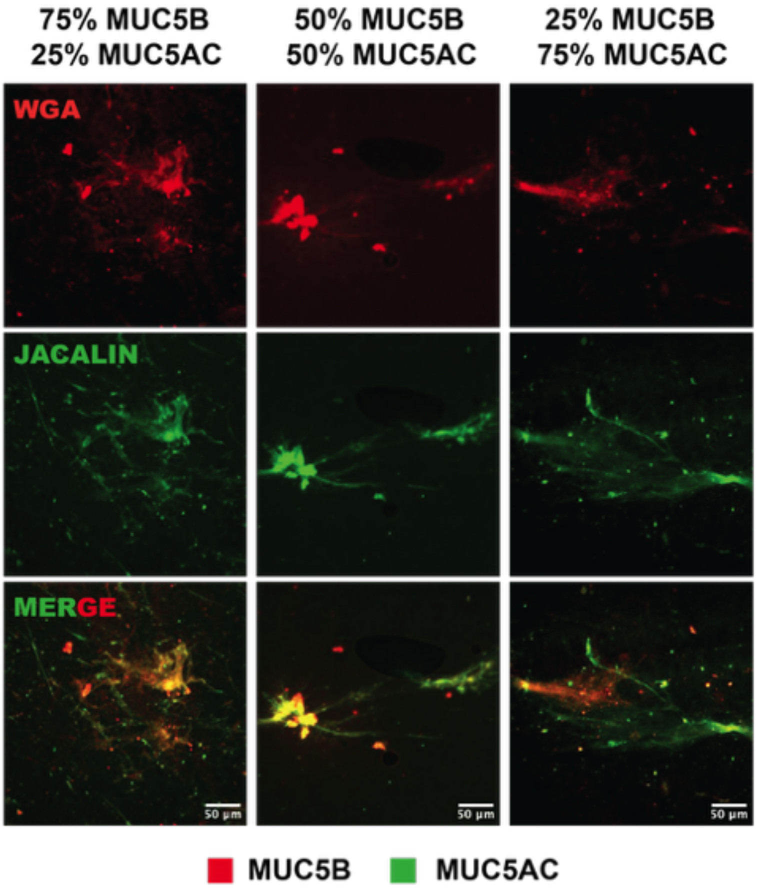


**Figure S2**. Lectin staining of mucus gels with varying MUC5B and MUC5AC mucin ratio with WGA (red) and Jacalin (green). Scale bar = 50 µm.

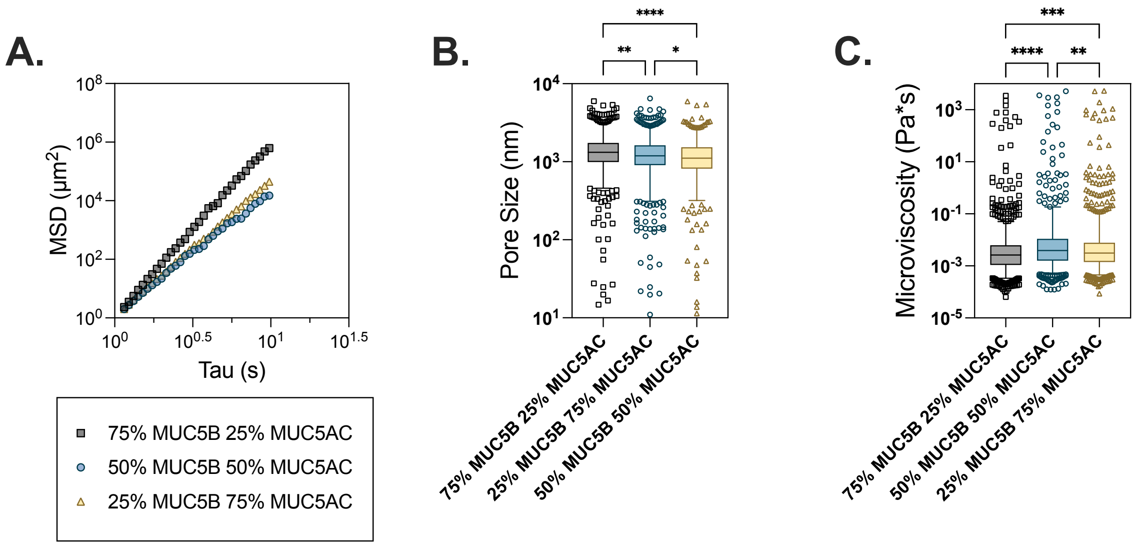


**Figure S3. Microrheological properties of mucus gels with varying MUC5B and MUC5AC ratios**. (**A**) Plot shows mean squared displacement vs time over 1 second. (**B-C**) Box-and-whisker plots of pore sizes and micro viscosity with varying mucin compositions from measured MSD values.

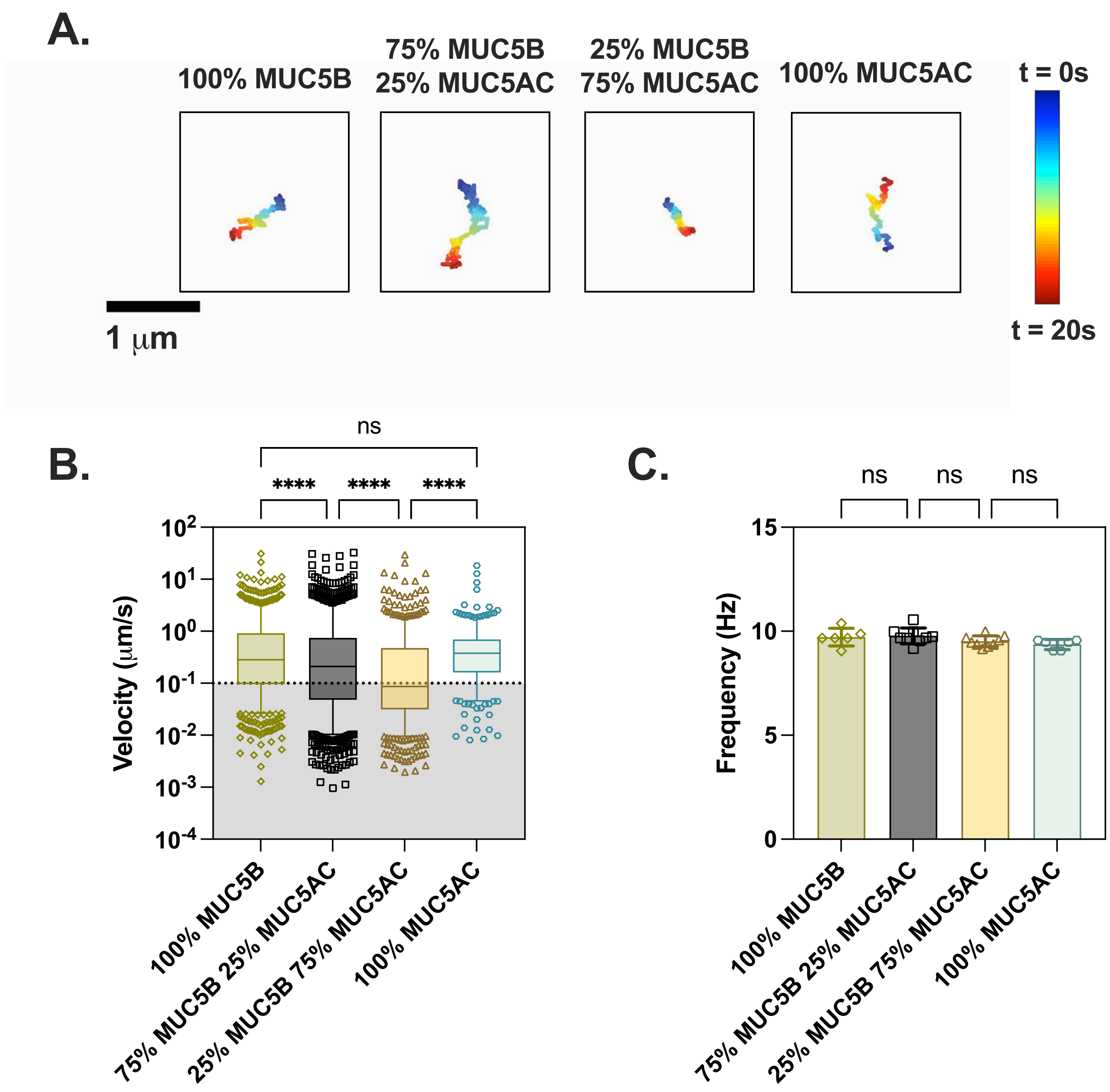


**Figure S4. Mucociliary transport in HAE cultures transplanted with mucus gels of varying mucin compositions.** (**A**) Representative trajectories of single beads in mucus gels with varying mucus compositions. Dark blue represents trajectories at 0 s, and dark red represents trajectories at 20 s. (**B**) Box-and-whisker plot representing velocities (μm/s) of all beads in mucus gels with varying mucus compositions. Shaded region indicates particles with velocities <0.1 μm/s. ****$p$<0.0001 by Kruskal-Wallis test with Dunn's correction. Each dot represents nanoparticle velocities from 5 videos per biological replicate (n=3). (**C**) Bar graph showing ciliary beat frequency (Hz) in mucus gels with varying mucus compositions. Each dot represents data from 1 video (n=3).

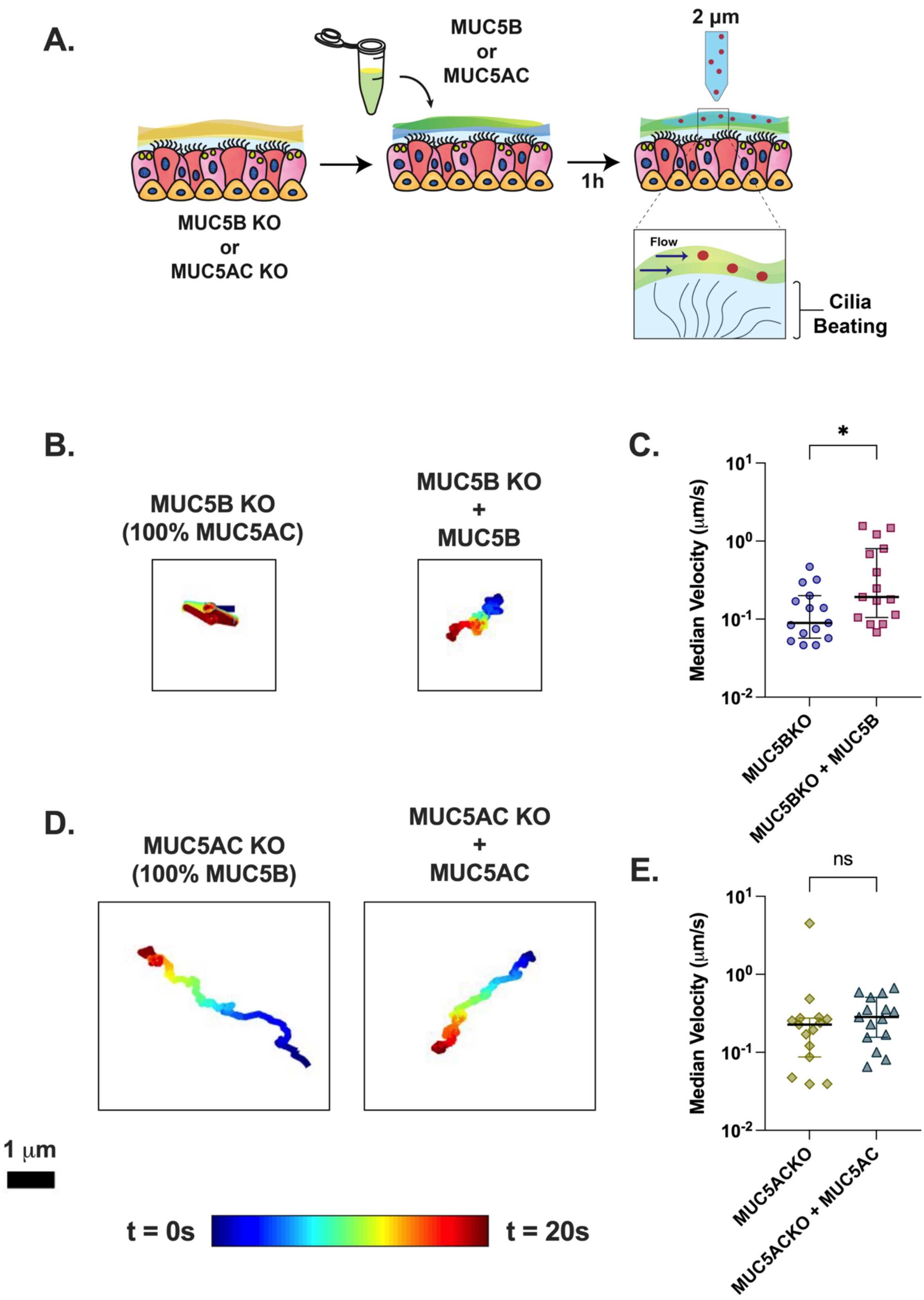


**Figure S5. Mucociliary transport rates after MUC5B and MUC5AC supplementation in KO cultures.** (A) Schematic illustration of MCT experiment design. HAE cultures are supplemented with the absent mucin on the apical surface. After 1 h of equilibration, 2 µm beads are applied to the apical surface, and the cultures are imaged to track the bead movement. (C) Representative trajectories of beads in MUC5B KO mucus and in MUC5B KO supplemented with MUC5B. (D) Scatter plot representing median velocities (µm/s) of beads in each video in MUC5B KO mucus and in MUC5B KO mucus supplemented with MUC5B. (E) Representative trajectories of beads in MUC5AC KO mucus and in MUC5B KO supplemented with MUC5AC. (F) Scatter plot representing median velocities (µm/s) of beads in each video in MUC5AC KO mucus and in MUC5AC KO mucus supplemented with MUC5AC. Each dot represents data from 1 video (n=3). $*p<0.05$ by one-way ANOVA.
